# Integrate Structural Analysis, Isoform Diversity, and Interferon-Inductive Propensity of ACE2 to Refine SARS-CoV2 Susceptibility Prediction in Vertebrates

**DOI:** 10.1101/2020.06.27.174961

**Authors:** Eric R. Sang, Yun Tian, Yuanying Gong, Laura C. Miller, Yongming Sang

**Affiliations:** Department of Agricultural and Environmental Sciences, College of Agriculture, Tennessee State University, 3500 John A. Merritt Boulevard, Nashville, TN, USA; Virus and Prion Diseases of Livestock Research Unit, National Animal Disease Center, USDA-ARS, Ames, IA, USA

**Keywords:** COVID-19, SARS-CoV2, Domestic animals, Angiotensin Converting Enzyme 2, Interferon signaling

## Abstract

The current new coronavirus disease (COVID-19) has caused globally near 0.4/6 million confirmed deaths/infected cases across more than 200 countries. As the etiological coronavirus (a.k.a. SARS-CoV2) may putatively have a bat origin, our understanding about its intermediate reservoir between bats and humans, especially its tropism in wild and domestic animals, are mostly unknown. This constitutes major concerns in public health for the current pandemics and potential zoonosis. Previous reports using structural analysis of the viral spike protein (S) binding its cell receptor of angiotensin-converting enzyme 2 (ACE2), indicate a broad SARS-CoV2 susceptibility in wild and particularly domestic animals. Through integration of key immunogenetic factors, including the existence of S-binding-void ACE2 isoforms and the disparity of ACE2 expression upon early innate immune response, we further refine the SARS-CoV2 susceptibility prediction to fit recent experimental validation. In addition to showing a broad susceptibility potential across mammalian species based on structural analysis, our results also reveal that domestic animals including dogs, pigs, cattle and goats may evolve ACE2-related immunogenetic diversity to restrict SARS-CoV2 infections. Thus, we propose that domestic animals may be unlikely to play a role as amplifying hosts unless the virus has further species-specific adaptation. These findings may relieve relevant public concerns regarding COVID-19-like risk in domestic animals, highlight virus-host coevolution, and evoke disease intervention through targeting ACE2 molecular diversity and interferon optimization.

## 1. Introduction

Erupting in China last December, the novel coronavirus disease (COVID-19) has become a worldwide pandemic and caused near 0.4 million confirmed deaths and 6 million infected cases across 200 countries by the end of May 2020 [1,2]. The etiological virus, designated as Severe Acute Respiratory Syndrome coronavirus 2 (SARS-CoV2) has been identified [3] and related to the viruses previously causing SARS or Middle East Respiratory Syndrome (MERS) in humans in 2003 and 2012, respectively [4]. These three human-pathogenic coronaviruses putatively evolve from bat coronaviruses, but have different animal tropisms and intermediate reservoirs before transmission to humans [4,5]. As civet cats and camels were retrospectively determined as reservoirs for SARS and MERS respectively, there is no conclusion about what animal species passing SARS-CoV2 to humans [4,5]. Investigations indicated that Canivora animals including raccoon dogs, red foxes, badgers and minks as well swine, at a less extent, are susceptible to SARS virus infections [6,7]. Although the viral nucleic acids and antibodies to MERS were detectable in multiple ruminant species including sheep, goat, and donkeys, the virus inoculation studies did not result in a productive infection for MERS disease in these domestic ruminants, nor in horses [8,9].

As a group of obligate pathogens, viruses need to engage cell receptors for entering cells and race with the host immunity for effective replication and spreading to initiate a productive infection [10]. In this context, the spike proteins protruding on the coronavirus surface are responsible for cell receptor binding and mediating viral entry [5–7]. For example, MERS-CoV adopts the dipeptidyl peptidase 4 (DPP4, a.k.a. CD26) and SARS-CoV uses angiotensin-converting enzyme 2 (ACE2) as primary receptors for cell attachment and entry [4–9]. Several groups have reported that SARS-CoV2 uses the same ACE2 receptor as SARS-CoV, but exerts higher receptor affinity to human ACE2, which may ascribe to the efficacy of SARS-CoV2 infection in humans [11,12]. After cell attachment via the receptor binding domain (RBD) in the N-terminal S1 region of the S protein, the C-terminal S2 region thus engages in membrane fusion. Further cleavage of S2 from S1 by a furin-like protease will release and prime the virus entering the recipient cells. Several furin-like proteases, especially a broadly expressed trans-membrane serine protease 2 (TMPRSS2), are adopted for priming SARS-CoV entry [11,12]. Compared with SARS-CoV, studies showed that SARS-CoV2 spike protein also evolutionarily obtains an additional furin-like proteinase cleavage site within the S1/S2 junction region for efficient release from the cell surface and entry into the cells [3,11–13].

Because TMPRSS2 is widely expressed, the tissue-specific expression of ACE2 has been shown to determine SARS-CoV2 cell tropism in humans [11,12]. Namely, human nasal secretory cells, type II pneumocytes, and absorptive enterocytes are ACE2-TMPRSS2 double positive and highly permissive to SARS-CoV2 infection [14, 15]. For cross-species animal tropism, the potential infectivity of SARS-CoV2 in both wild and domestic animals raises a big public health concern after the prevalence of SARS-CoV2 infections in humans [16,17]. This concern involves two aspects: (1) screening to identify the animal species that serve as a virus reservoir originally passing SARS-CoV2 to humans; and (2) the existing risk of infected people passing the virus to animals, particularly domestic species, thus potentially amplifying the zoonotic cycle to worsen SARS-CoV2 evolution and prevalence [16,17]. By diagnosis of animals in close contact with COVID-19 patients or screening of animal samples in some COVID-19 epidemic zones, studies detected that domestic cats and dogs could be virally or serologically positive for SARS-CoV2 [18–24], as was a reported infection in a zoo tiger [25]. Using controlled experimental infection of human SARS-CoV2 isolates, several studies demonstrated that ferrets, hamsters, domestic cats and some non-human primate species are susceptible to human SARS-CoV2 strains [18–25]. Obviously, it is impractical to test SARS-CoV2 susceptibility experimentally in all animal species. By adoption of a structural simulation based on published structures of the viral S-RBD/ACE2 complex, studies have predicted a broad spectrum of vertebrate species with high potential for SARS-CoV2 susceptibility, which, if true, entails unexpected risks in both public and animal health, and warrants further critical evaluation [26–28].

ACE2 is a key enzyme catalyzing angiotensin (AGT) further conversion into numeral active forms of AGT1-9, which are hormonal mediators in the body’s renin-angiotensin system (RAS) [29,30]. Thus, ACE2 plays a regulatory role in the blood volume/pressure, body fluid balance, sodium and water retention, as well as immune effects on apoptosis, inflammation, and generation of reactive oxygen species (ROS) [29,30]. In this line, the expression of ACE2 is also inter-regulated by immune mediators pertinent to its systemic function. Multiple physio-pathological factors, including pathogenic inflammation, influence on RAS through action on ACE2 expression [29–31]. Interferon (IFN) response, especially that mediated by type I and type III IFNs, comprises a frontline of antiviral immunity to restrict viral spreading from the initial infection sites, and therefore primarily determines if a viral exposure becomes controlled or a productive infection [32]. Several recent studies revealed that human *ACE2* gene behaves like an interferon-stimulated gene (ISG) and is stimulated by a viral infection and IFN treatment; however, mouse *ACE2* gene is not [15,33,34]. Therefore, to determine the cell tropism and animal susceptibility to SARS-CoV2, the cross-species *ACE2* genetic and especially epigenetic diversity in regulation of ACE2 expression and functionality should be evaluated [26–34]. In this study, through integration of structural analysis and key immunogenetic factors that show species-dependent differences, we critically refine the SARS-CoV2 susceptibility prediction to fit recent experimental validation [16–25]. Along with showing a broad susceptibility potential across mammalian species based on structural analysis [26–28], our results further reveal that domestic animals including dogs, pigs, cattle and goats may evolve previously unexamined immunogenetic diversity to restrict SARS-CoV2 infections.

## 2. Materials and Methods

### Protein and promoter sequence extraction and alignment

The amino acid sequences of ACE2 proteins and DNA sequences of the proximal promoters of each *ACE2* genes were extracted from NCBI Gene and relevant databases (https://www.ncbi.nlm.nih.gov/gene). *ACE2* genes and corresponding transcripts have been well annotated in most representative vertebrate species. In most cases, the annotations were double verified through the same Gene entries at Ensembl (https://www.ensembl.org). The protein sequences were collected from all non-redundant transcript variants and further verified for expression using relevant RNA-Seq data (NCBI GEO profiles). The proximal promoter region spans ~2.5 kb before the predicted transcription (or translation) start site (TSS) of ACE2 or other genes. The protein and DNA sequences were aligned using the multiple sequence alignment tools of ClustalW or Muscle through an EMBL-EBI port (https://www.ebi.ac.uk/). Other sequence management was conducted using programs at the Sequence Manipulation Suite (http://www.bioinformatics.org). Sequence alignments were visualized using Jalview (http://www.jalview.org) and MEGAx (https://www.megasoftware.net). Sequence similarity calculations and plotting were done using SDT1.2 (http://web.cbio.uct.ac.za/~brejnev). Other than indicated, all programs were run with default parameters.

### Phylogenic analysis

The phylogenic analysis and tree visualization were performed using MEGAx and an online program, EvoView. The evolutionary history was inferred using the Neighbor-Joining method. Percentage of replicate trees in which the associated taxa clustered together in the bootstrap test (1,000 replicates) was also performed. The evolutionary distances were computed using the p-distance method and in units of the number of amino acid differences per site. Other than indicated, all programs were run with default parameters as the programs suggested.

### Structural simulation and analysis

The structure files of human ACE2 protein and its interaction with SARS-CoV2 S-RBD were extracted from the Protein Data Bank under the files of 6M17 and 6M0J. The residual mutation and structure simulation were performed using UCSF Chimera and Pymol available at https://www.cgl.ucsf.edu/chimera/ and https://pymol.org/, respectively. Structural visualization were using Pymol. The binding affinity energy (ΔG), dissociation constant (Kd) and interfacial contacts between S-RBD and each ACE2 were calculated using an PRODIGY algorithm at https://bianca.science.uu.nl/prodigy/.

### Profiling transcription factor binding sites in ACE2 promoters and PWM scoring

The regulatory elements (and pertinent binding factors) in the ~2.5 kb proximal promoter regions was examined against both human/animal TFD Database using a program Nsite (Version 5.2013, at http://www.softberry.com). The mean position weight matrix (PWM) of key cis-elements in the proximal promoters were calculated using PWM tools through https://ccg.epfl.ch/cgi-bin/pwmtools, and the binding motif matrices of examined TFs were extracted from JASPAR Core 2018 vertebrates (http://jaspar.genereg.net/).

### RNA-Seq and data analysis

For expression confirmation, several sets of RNA-Seq data from NCBI Gene databases, and one of ours generated from porcine alveolar macrophages (BioProject with an accession number of SRP033717), were analyzed for verification of the differential expression of *ACE2* genes in most annotated animal species. Especially, the expression of porcine ACE2 isoforms and relevant other genes in the porcine lung macrophage datasets. Significantly differentially expressed genes (DEGs) between two treatments were called using an edgeR package and visualized using heatmaps or bar charts as previously described [58].

## 3. Results and Discussion

### 3.1. Vertebrate ACE2 orthologs share an functional constraint but experience intra-species diversification in livestock with unknown selective pressure

Sequence comparison among ACE2 orthologs across 30 representative vertebrate species shows a pairwise identity range at 57-85% (Fig. 1A and Supplemental Fig. S1 and Excel Sheet), which is 15-27% higher than the average value generated through a similarity analysis at 30-70% on gene orthologs at a genome-wide scale [35]. This indicates that ACE2 exerts a similar and basic function cross-species, consistent with its systemic and regulatory role as a key enzyme in RAS, an essential regulatory axis underlying the body circulatory and execratory systems in vertebrates [29–31]. A comparison of evolutionary rates of major genes within RAS including angiotensinogen (AGT), ACE, and several receptors of the processed angiotensin hormones showed that ACE2 actually evolves slightly faster than ACE [36, and unpublished data]. This implies that ACE2 may bear pressure for RAS adapting evolution per a species-dependent physiological and pathological requirement [29–31]. This evolutionary adaptability of *ACE2* genes is demonstrated by the existence of numerical genetic polymorphisms [37] and several transcript isoforms particularly in humans and major livestock species (Fig. 1B and Supplemental Fig. S1 and Excel Sheet). We identified (and verified by RNA-Seq annotation) four transcripts of ACE2 isoforms in humans (Fig. 1B) that primarily differ in the C-terminal 50 residues within the collectrin domain. Particularly, 1-2 short ACE2 isoforms were identified in dogs, pigs, cattle, and goats in addition to the longer ACE2 consensus to the human’s (designated as –S or –L, respectively after the animal common names in Fig. 1B and thereafter). These livestock ACE2-S isoforms have a 70-130 residual truncation at their N-terminal peptidase domains, which also span the region interacting with SARS-CoV spike protein. The selective mechanisms driving the evolution of these short ACE2 isoforms in livestock are unknown, but may relate to previous pathogenic exposure or unprecedented physio-pathological pressure. To support this reasoning, short ACE2 isoforms are detected in both domestic *Bos taurus* and hybrid cattle, but not in the wild buffalo and bison; and ACE2 isoforms from each species are generally paralogous and sister each other within a clade in the phylogenic tree (Fig. 1B).

**Figure 1:**
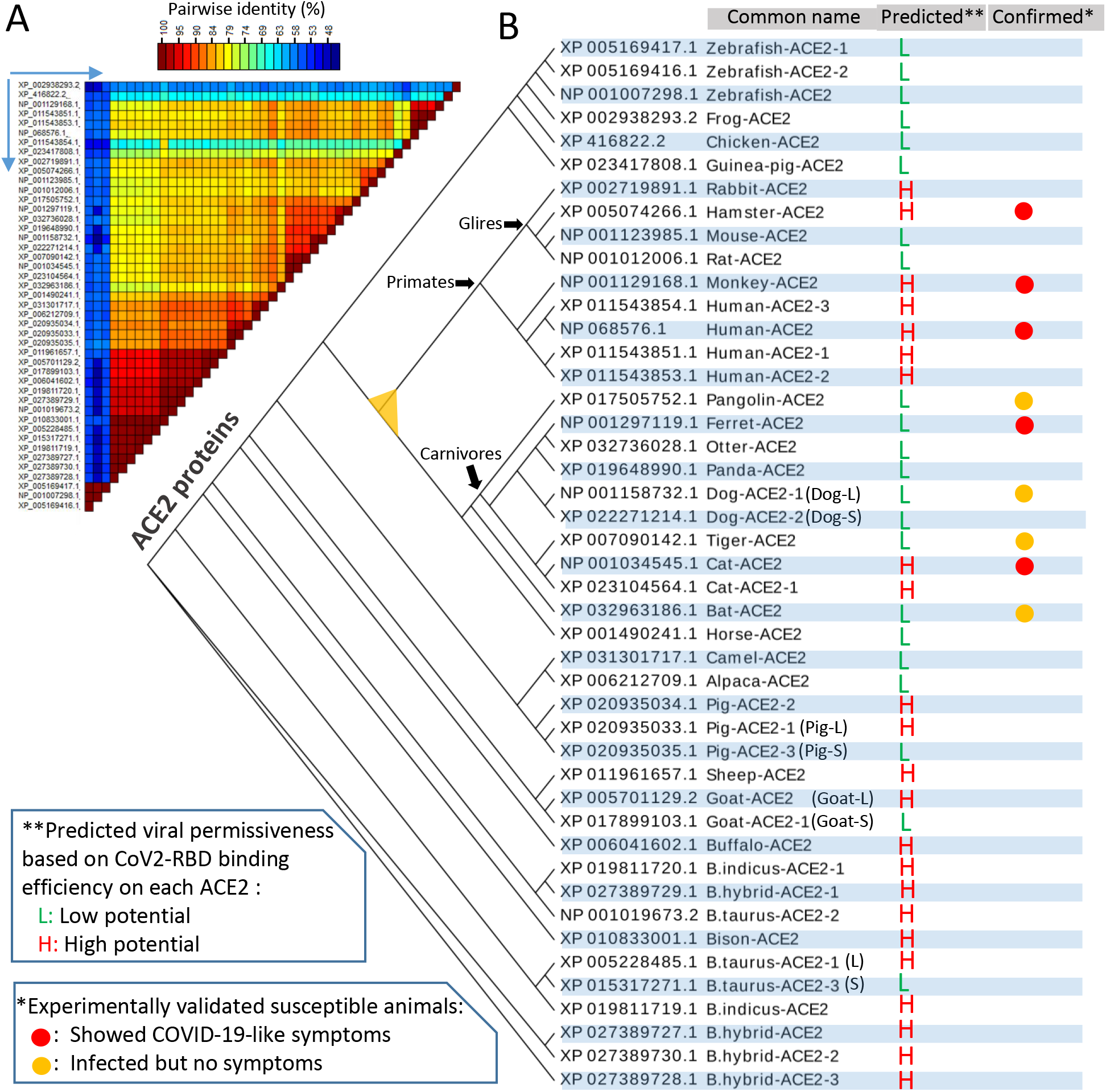
Incongruence of predicted SARS-CoV2 susceptibility to experimental data in selected major livestock and reference animal species. Protein sequences of angiotensin converting enzyme 2 (ACE2) orthologs in different species, and transcript variants identified within some species, were extracted from NCBI (and Ensembl) Gene databases and verified using relevant RNA-Seq expression. **(A)** Sequence alignment was performed with ClustalW and pairwise identity (%) was calculated and visualized using SDT1.2. The reciprocal proteins for comparison on the horizontal are the same order as the vertical. Only NCBI Accession numbers are listed, see the common names in (B). **(B)** The phylogenic tree of major identified ACE2 orthologs/variants from different species was built with a Neighbor-joining approach and visualized using an EvoView program under default parameter setting. The prediction of SARS-CoV2 susceptibility is based on the sequence similarity of each ACE2 to human ACE2 in the S-RBD binding region and simulated using a published human ACE2-RBD structure (6M0J) and refers to two recent publications using similar procedures but different structural models [27,28]. Compared with the currently available experimental data, incongruence of the predicted SARS-CoV2 susceptibility is clearly demonstrated in pangolin, ferret, tiger, cat and horseshoe bat, indicating that some other factors besides ACE2-RBD affinity should be considered.

### 3.2. Incongruence of predicted SARS-CoV2 susceptibility to experimental data in selected major livestock and reference animal species

Phylogenic analysis of vertebrate ACE2 orthologs/paralogs reveals a general relationship aligning to the animal cladistics (Fig. 1B). In this context, homologs from the fish, frog and chicken conform to a primitive clade. All ungulate homologs form into parallel clades next to each other. The homologs from the glires, primates and carnivores cluster into a big clade (marked with yellow triangle node), which contains all the SARS-CoV2 susceptible species that have been verified via natural exposure or experimental infections (Fig. 1B, marked with red/orange circles). We examined and merged several previous studies about the prediction of SARS-CoV2 susceptibility in vertebrates based on the simulated structural analysis of S-RBD-ACE2 complex [26–28]. As numerous vertebrate species were predicted to be high or low potential (Fig. 1B, labeled as red H or green L) for SARS-CoV2 susceptibility, incongruence between the predicted SARS-CoV2 susceptibility and infected validation is apparent in pangolin, ferret, tiger, cat and horseshoe bat, indicating that some other factors besides ACE2-RBD affinity should be considered [15,32–34]. We, therefore, refined the prediction matrix to include the RBD-binding evasion of some ACE2 orthologs identified in major livestock species and the interferon-stimulated ACE2 expression underlying SARS-CoV2 infections [15,32–34].

### 3.3. A broad SARS-CoV2 susceptibility potential based on structural analysis

Several recent studies have elegantly demonstrated the structural interaction of the viral S protein or its RBD in complex with human ACE2 receptor [38,39]. Showing that the contacting residues at the RBD/ACE2 interface (Fig. 2A) involve at least 19 residues in ACE2 (Fig. 2B, listed in the Table cells and referred to the aligned residual positions in human ACE2) and 10 residues in the SARS-CoV2 RBD (Fig. 2B, blue circles with residue labels above the Table) [28,38,39]. The cross-species residual identity (%) of these interacting residues in ACE2 are dispersed in a broader range (32-100%) than the whole ACE2 sequence identity rate at 57-85% [35], indicating a faster evolution rate of this virus-interacting region. Notably, the S-binding region spans a large part of the N-terminal peptidase domain and S-binding may competitively block a majority of active sites of the enzyme (Fig. 2C).

**Figure 2:**
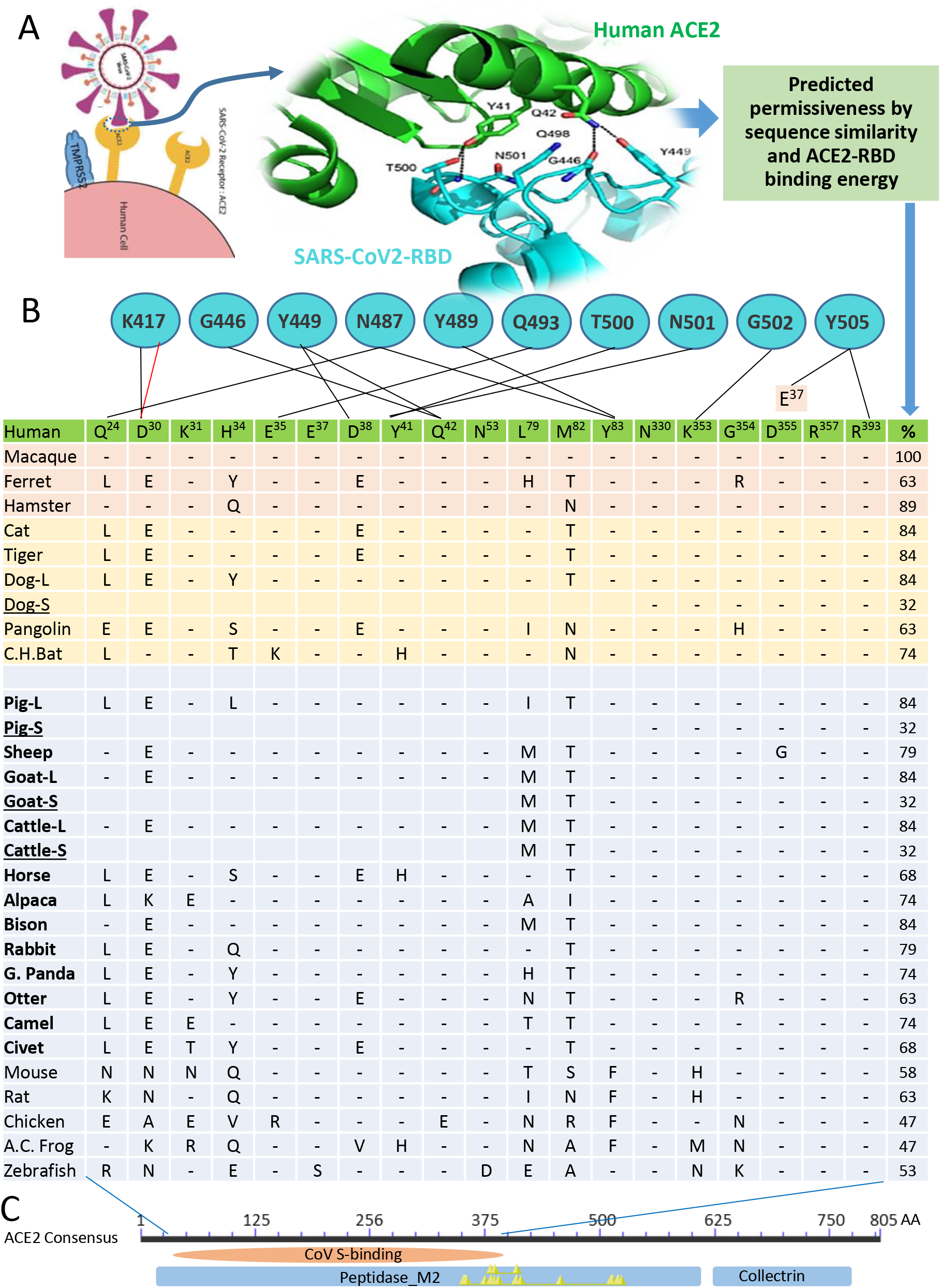
Prediction of SARS-CoV2 susceptibility in major livestock species based on the conservation of key interacting residues and binding capacity between the viral spike (S) protein on the host ACE2 receptor. **(A)** SARS-CoV-2 uses the cell receptor, angiotensin-converting enzyme 2 (ACE2) for entry and the serine protease TMPRSS2 and furin for S protein priming. **(B)** As TMPRSS2 is broadly expressed and active with a furin-like cleavage activity, the affinity adaption of the S receptor binding domain (RBD) and ACE2 receptor determines the viral permissiveness. The contacting residues of human ACE2 (a distance cutoff 4.5 Å) at the SARS-CoV-2 RBD/ACE2 interfaces are shown, and the contacting network involves at least 19 residues in ACE2 (listed in the Table cells and refer to the aligned residual positions in human ACE2) and 10 residues in the SARS-CoV-2 RBD (blue circles with residue labels), which are listed and connected with black lines (indicating hydrogen bonds) and red line (represents salt-bridge interaction). The cross-species residual identity (%) of these interacting residues in ACE2 are listed in a broad range (32-100%) [26–28]. **(C)** We also detected several short ACE2 isoforms (underlined) in the domestic animals including dog, pig, goat and cattle, which have a N-terminal truncation spanning 10-13 key residues in the contacting network to S-RBD but keeping the enzyme active sites (indicated by Yellow triangles), thus resulting in little engagement by the viral S protein and predicting an unexpected evolutionary advantage for relieving potential COVID-19 risk caused by the viral engagement and functional distortion on the classical long ACE2 isoforms in these animal species. The NCBI Accession Numbers of the ACE2 orthologs are listed as in Fig. 1.

Using a similar structural analysis procedure [27,28], we modeled the ACE2 structures of animal species of interest and simulated their interaction with SARS-CoV2 S-RBD based on a published RBD-human ACE2 structure (Protein Data Bank File 6M0J) [38]. Fig. 3 demonstrates the S-RBD interaction with the simulated structures of ACE2 long isoforms from the dog, pig and cattle, respectively. The major changes of the RBD-ACE2 interacting interfaces are from the residual exchanges in ACE2 from other species compared with human ACE2 (Fig. 2B-2D, highlighted in red). In addition, the exchange of N90T (in pigs) and N322Y (in cattle and sheep) would destroy the N-glycosylation site in human ACE2. ACE2 from goat (Supplement Fig. S1) exhibits identical amino acid exchanges as in cattle in the RBD-ACE2 interfacial contacts. In contrast,when compared with human ACE2, ACE2 from cats (Supplement Fig. S1) conserves all relevant glycosylation sites in human ACE2 [28,38]. We also calculated the interfacial contacts using parameters of protein-protein interaction including the predictable binding affinity energy (ΔG), dissociation constant (Kd) and number of different interfacial contacts within the S-RBD and ACE2 contact. Although the exact numbers may differ from previous reports [38], they provide a very comparable matrix generated using the same algorithm (Fig. 3E) [40]. Data show that the ACE2 of most domestic animals, including that from mouse and rat (species known to be unsusceptible to human SARS-CoV2), have a binding affinity (ΔG) at −11.2 to −12.8 kcal/mol. This is within the binding affinity range (11.2-12.9 kcal/mol) between the RBD and the ACE2 from known susceptible species (Fig. 3E, underlined in the left part of the table). This indicates that other factors, conceivably from genetic divergence and/or natural immunity, also contribute to SARS-CoV2 susceptibility in animal species. Therefore, an effective prediction matrix should include the critical immunogenetic factors to further determine virus susceptibility in addition to the sequence/structural similarity of ACE2 receptors (Fig. 1 and Fig. S1) [15,34,37].

**Figure 3:**
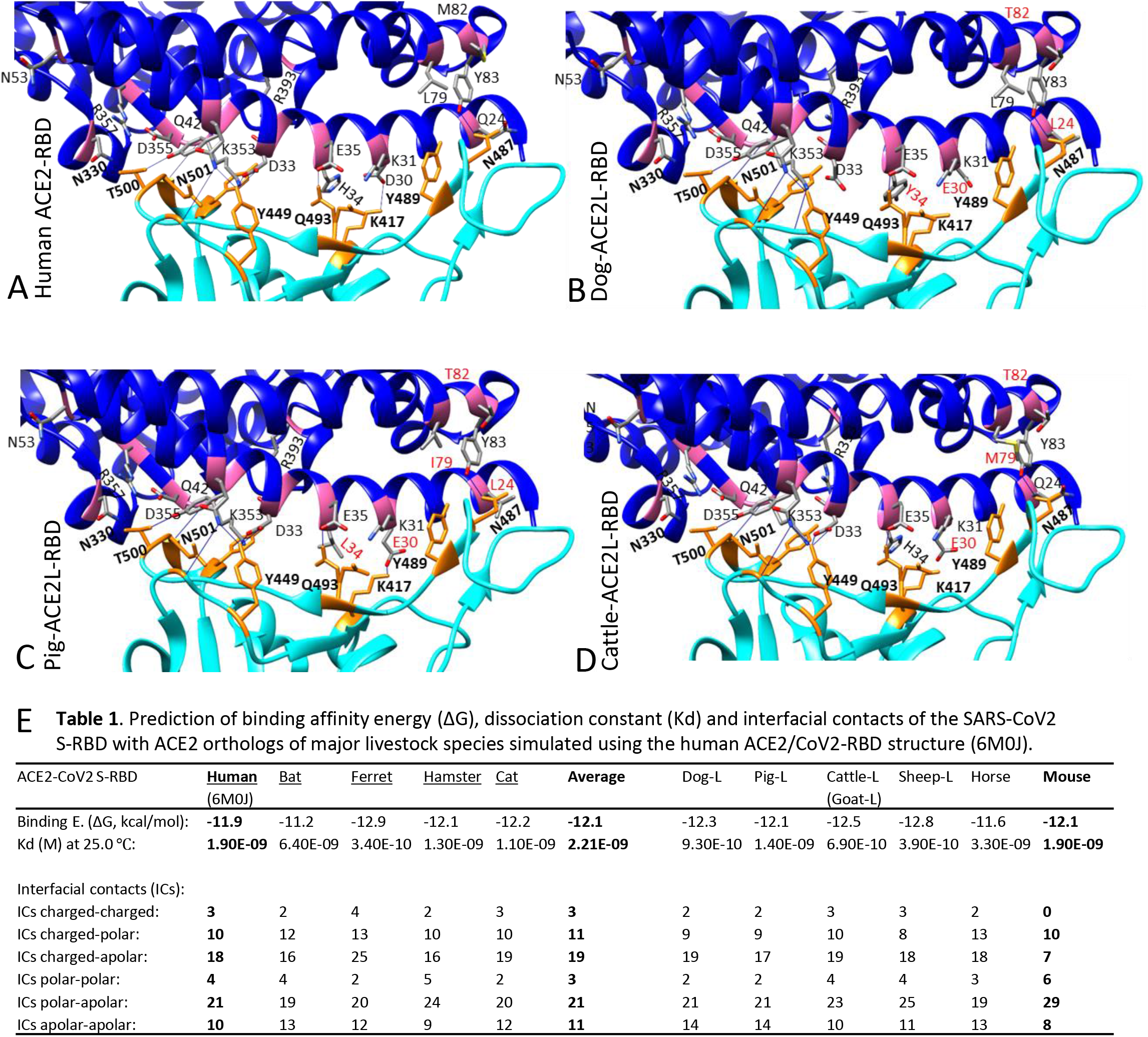
Binding capacity of SARS-CoV2 spike protein (S) and its cell receptors (ACE2) from different animal species. **(A)** Structure of the receptor-binding domain (RBD) of S from SARS-CoV2 (cyan) bound to human angiotensin-converting enzyme 2 (ACE2) (blue). Most residues involved in binding are highlighted as magenta (ACE2) or orange (S) sticks and labeled as one-letter amino-acid codes plus residual numbers in **bold** or regular font respectively for S or ACE2 residues. The dotted/blue lines indicate intermolecular salt bridge or hydrogen bonds between interacting residues (generated and visualized with UCSF Chimera and Pymol from Protein Data Bank File 6M0J). **(B) to (D)** RBD interaction with the simulated structures of ACE2 long isoforms from the dog, pig and cattle, respectively. Amino acid exchanges in ACE2 from another species compared with human ACE2 are highlighted in red. **(E)** Prediction of binding affinity energy (ΔG), dissociation constant (Kd) and interfacial contacts of the SARS-CoV2 S-RBD with ACE2 orthologs of major livestock species. Most domestic animals ACE2 including that from mouse and rat (species known not to be susceptible to SARS-CoV2) have a binding affinity (ΔG) at −11.2 to −12.8 kcal/mol that is within the range (11.2-12.9 kcal/mol) between the RBD and the ACE2 from the known susceptible species (underlined in the left part of the table), indicating that some other factors, especially those from genetic divergence and natural immunity, contribute to the SARS-CoV2 susceptibility of different animal species.

### 3.4. Identification of livestock short ACE2 isoforms that likely evade the binding by SARS-CoV2 Spike protein

We detected several short ACE2 isoforms in the domestic animals including dog, pig, goat and cattle that have an N-terminal truncation spanning 10-13 key residues in the contacting network to S-RBD but retain the enzyme active sites (Fig. 4A). Most of the splicing isoforms of *ACE2* genes, such as in zebrafish, cats and humans, share a common proximal promoter and encode ACE2 proteins containing all 19 key RBD-interacting residues [38,39]. However, these short ACE2-S isoforms in domestic animals truncate for 71 (Cattle/Goat ACE2-S) or 132 (Dog/Pig ACE2-S) residues at their N-termini compared with the long ACE2 isoforms in the same species (Fig. 1 and Fig. S1). Therefore, these short ACE2 isoforms destroy 10-13 key residues in the contacting network to S-RBD but likely retain ACE2 enzymatic function in RAS. Paired structural comparison between the human ACE2 structure (extracted from 6M17) with each simulated ACE2-S structure from the pig, dog, and cattle/goat, reveals that all these ACE2-S orthologs from domestic animals, particularly the porcine one, show high structural similarity to the human ACE2 except for the N-terminal truncations (Fig. 4B-4D). This indicates that these short ACE2 isoforms in domestic animals have little chance to be engaged by the viral S-binding, and predict an unexpected evolutionary advantage to allay potential COVID-19 risk resulting from viral engagement and functional distortion on the classical long ACE2 isoforms in these animal species [37,41].

**Figure 4:**
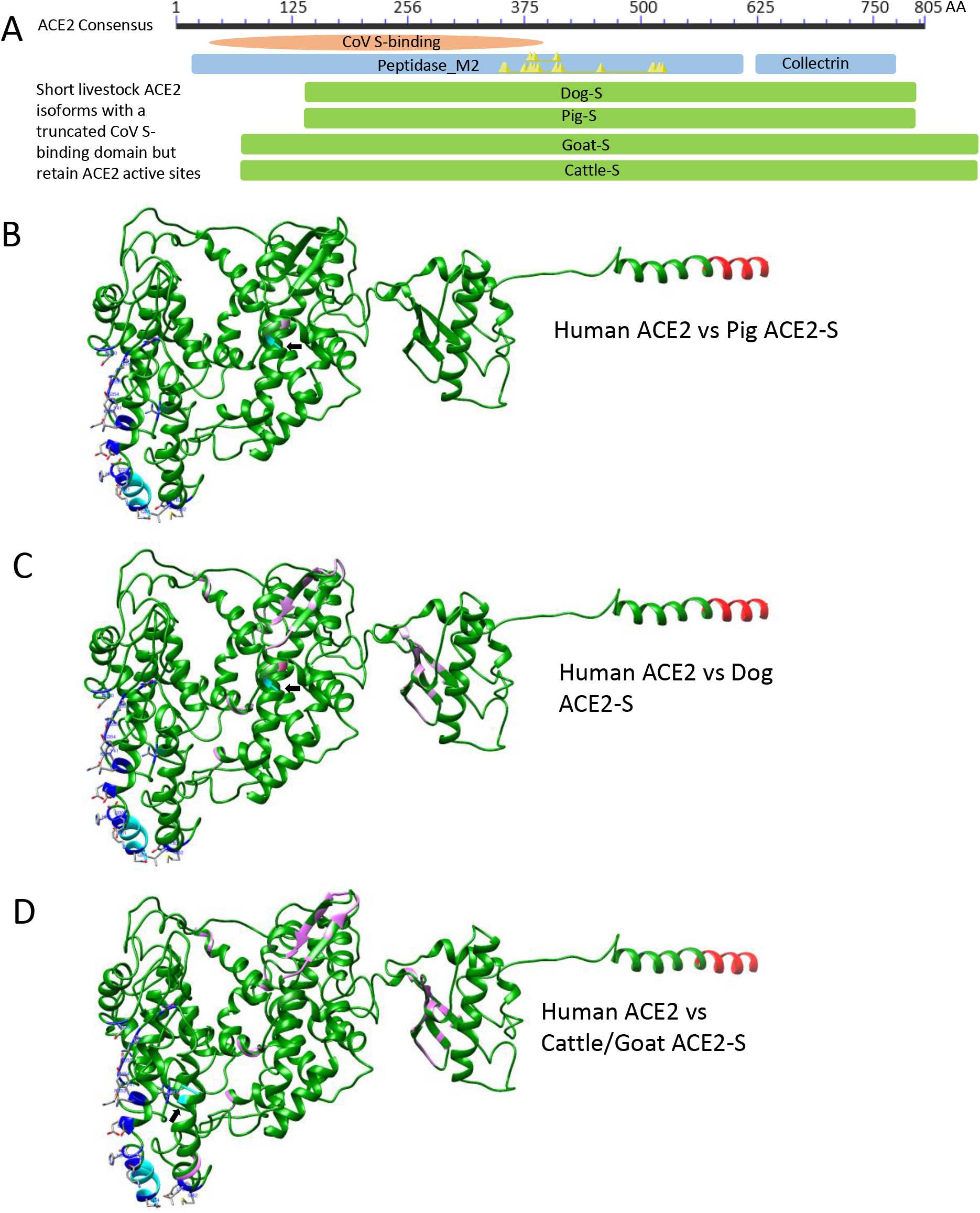
Detection of several short ACE2 isoforms (ACE2-S) in the domestic animals including dog, pig, goat and cattle. **(A)** In contrast to most splicing isoforms such as in cats and humans, which share a common proximal promoter and encode ACE2 proteins with similar sequences containing all 19 key RBD-interacting residues, these short ACE2-S isoforms in domestic animals truncate for 71 (Cattle/Goat ACE2-S) or 132 (Dog/Pig ACE2-S) residues at their N-termini compared with human ACE2 or the long ACE2 isoforms in these species, thus destroying 10-13 key residues in the contacting network to S-RBD but retaining all enzyme active sites (Yellow triangles in the blue ACE2 domain bar). This results in little chance to be engaged by the viral S protein binding and predicts an unexpected evolutionary advantage to relieve potential COVID-19 risk caused by the viral engagement and functional distortion on the classical long ACE2 isoforms in these animal species. **(B), (C)** and **(D)** Paired structural comparison between the human ACE2 structure (6M17) with each simulated ACE2-S structure from pig (B), dog (C) and cattle/goat (D). Human ACE2 structure are in green, and each compared animal ACE2-S structure in magenta. The N-terminal residues of both compared structures are in cyan (arrows indicating N-termini of the ACE2-S isoforms) and shared C-termini are in red. The 19 key S-interacting residues in human ACE2 are shown in blue sticks. In general, all ACE2-S orthologs, particular the porcine, show high structural similarity to the human ACE2 except the N-terminal truncations.

### 3.5. ACE2 *gene*s in vertebrates obtain different propensity in response to viral infection and interferon stimulation

SARS-CoV2 infection induces a weak IFN response but a production of a high amount of inflammatory cytokines including interleukin (IL)-6 and chemokine CXCL10 in most severe COVID-19 patients [42–45]. Studies of SARS and MERS showed that these pathogenic coronaviruses share similar viral antagonisms, including the endoribonuclease (EndoU) encoded by nonstructural protein 15 (nsp15), which directly blunts cell receptors responding to viral dsRNA and in turn weaken the acute antiviral response [46]. Several recent studies revealed that SARS-CoV2 seems more cunning in not only evading or antagonizing but also in exploiting the IFN response for efficient cell attachment [15,42,43,46]. As a key enzyme in RAS, the expression of *ACE2* gene has been primarily investigated for physiological response to circulatory regulations, and a response to pathological inflammation is also expected [29–31]. However, the expression of the *ACE2* gene was highly responsive to both viral infection and host IFN response, i.e. human *ACE2* gene seems an unstudied IFN-stimulated gene (ISG) [15,33]. Surprisingly, the ISG propensity of *ACE2* genes is species-dependent, for example: the mouse *ACE2* gene is less IFN responsive which may partly explain the mouse insusceptibility to SARS-CoV2 infection [15].

To categorize the different IFN-inductive propensity of *ACE2* genes in vertebrates, particularly in major livestock species, we profiled the regulatory cis-elements and relevant transcription factors in the proximal promoter regions of each *ACE2* genes (2.5 Kb before TSS or ATG). Figure 5 illustrates major regulatory cis-elements located in *ACE2* genes from major livestock animals and several reference animal species. Data show that animal *ACE2* gene promoters are evolutionally different in containing IFN- or virus-stimulated response elements (ISRE, PRDI, IFRs, and/or STAT1/3 factors) and cis-elements responsive to pro-inflammatory mediators. All these cis-elements recruit corresponding transcription factors (TF) to mediate differential ACE2 responses to antiviral IFNs and inflammation that is associated with COVID-19 disease [2,3,47]. We discover that *ACE2* genes obtain species-different ISG propensity responsive to IFN and inflammatory stimuli. In most (if not all) of the SARS-CoV2 susceptible species the *ACE2* genes obtained the IFN response between the typical robust and tunable IFN-stimulated genes (ISG) [48]. In general, the robust ISGs (ISG15 is an example here) are stimulated in the acute phase of viral infection and play a more antiviral role; in contrast, the later responsive tunable ISGs (IRF1 is an example) contribute more to anti-proliferation of IFN activity [48]. In addition, unlike the promoter of the short ACE2 isoforms in cattle and goats, which share most common promoter regions with their paralogous long isoforms, the short ACE2 isoforms of dogs (Dog-S) and pigs (Pig-S) have distinct proximal promoter regions (and different IFN responsivity) to the paralogous long ACE2 isoforms (Fig. 5 and Fig 6). Results indicate that the short ACE2 isoforms in pigs and dogs diversify from their long paralogs at both the levels of genetic coding and epigenetic regulation to adapt to some evolutionary pressure, such as that from pathogenic interaction (Fig. 7) [37,49].

**Figure 5.**
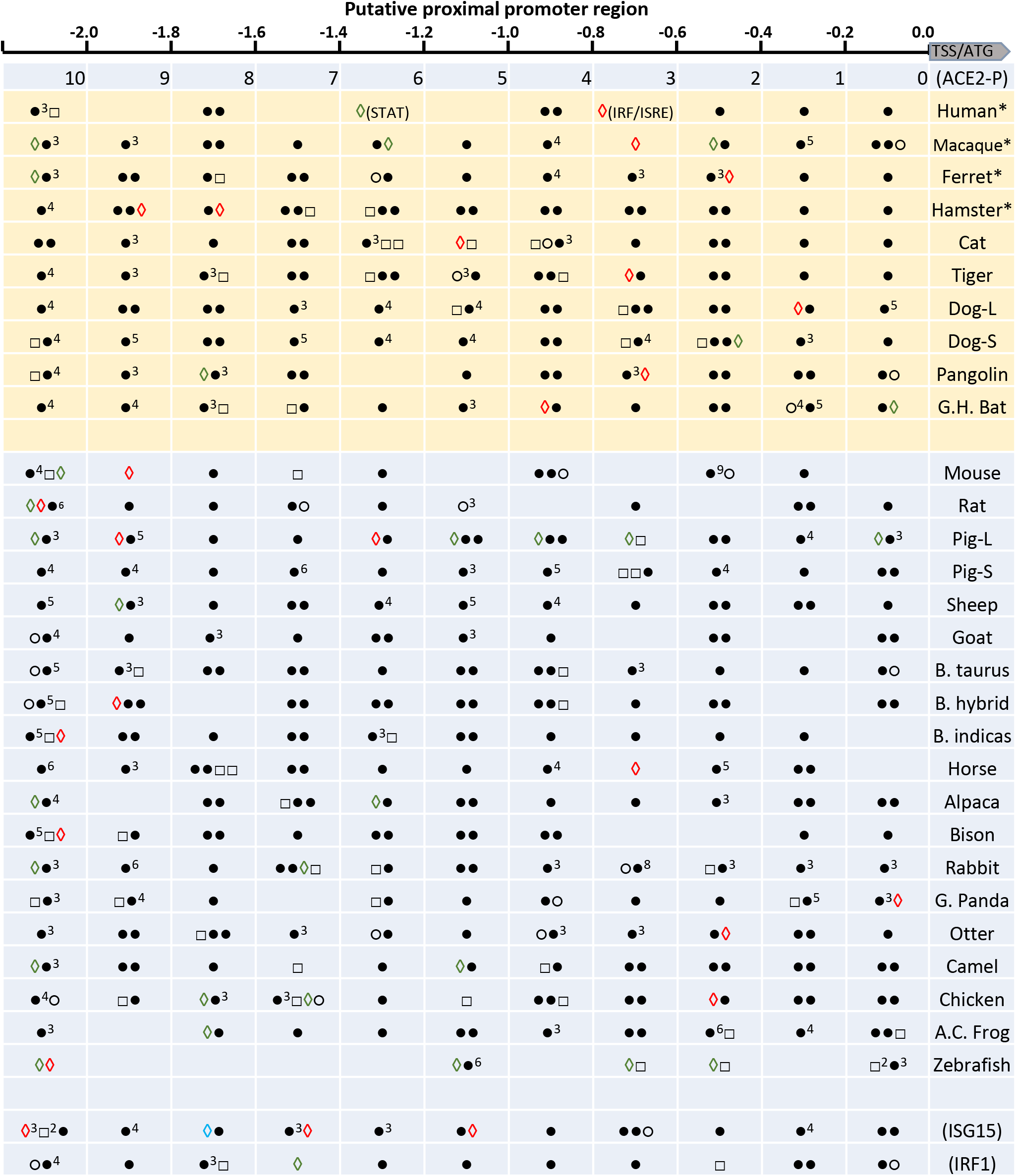
Categorizing *ACE2* genes based on regulatory cis-elements predicted in their proximal promoter regions (<2 Kb before TSS or ATG). The regulatory elements (and pertinent binding factors) in the ~2 kb proximal promoter regions were examined against both human/animal TFD Database using a program Nsite (Version 5.2013, at http://www.softberry.com), including *ACE2* genes identified in major livestock animals and several reference animal species. Data show that animal *ACE2* gene promoters are evolutionally different in containing IFN- or virus-stimulated response elements (ISRE, PRDI, IFRs, and/or STAT1/3 factors) and cis-elements responsive to pro-inflammatory mediators, which mediate different ACE2 responses to antiviral interferons (IFNs) and inflammation associated with COVID-19 disease. *Legend:* ∘, GATA-1 regulating constitutive expression;, acute 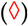 or secondary 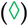 IFN-stimulated response element (ISRE) and PRDI that interact with IRF, ISGF3 and STAT factors, respectively; □, cis-elements interacting with factors to mediate immune/inflammatory responses including C/EBP, NF-kB, NF-IL6, and p53; •, cis-elements reacting with other factors significant in other developmental/physiological responses. The promoter features of two typical human interferon-stimulated genes (ISG), the robust ISG15 and tunable IRF1 are shown as references to indicate that *ACE2* genes obtain species-different ISG propensity responsive to IFN and inflammatory stimuli.

**Figure 6.**
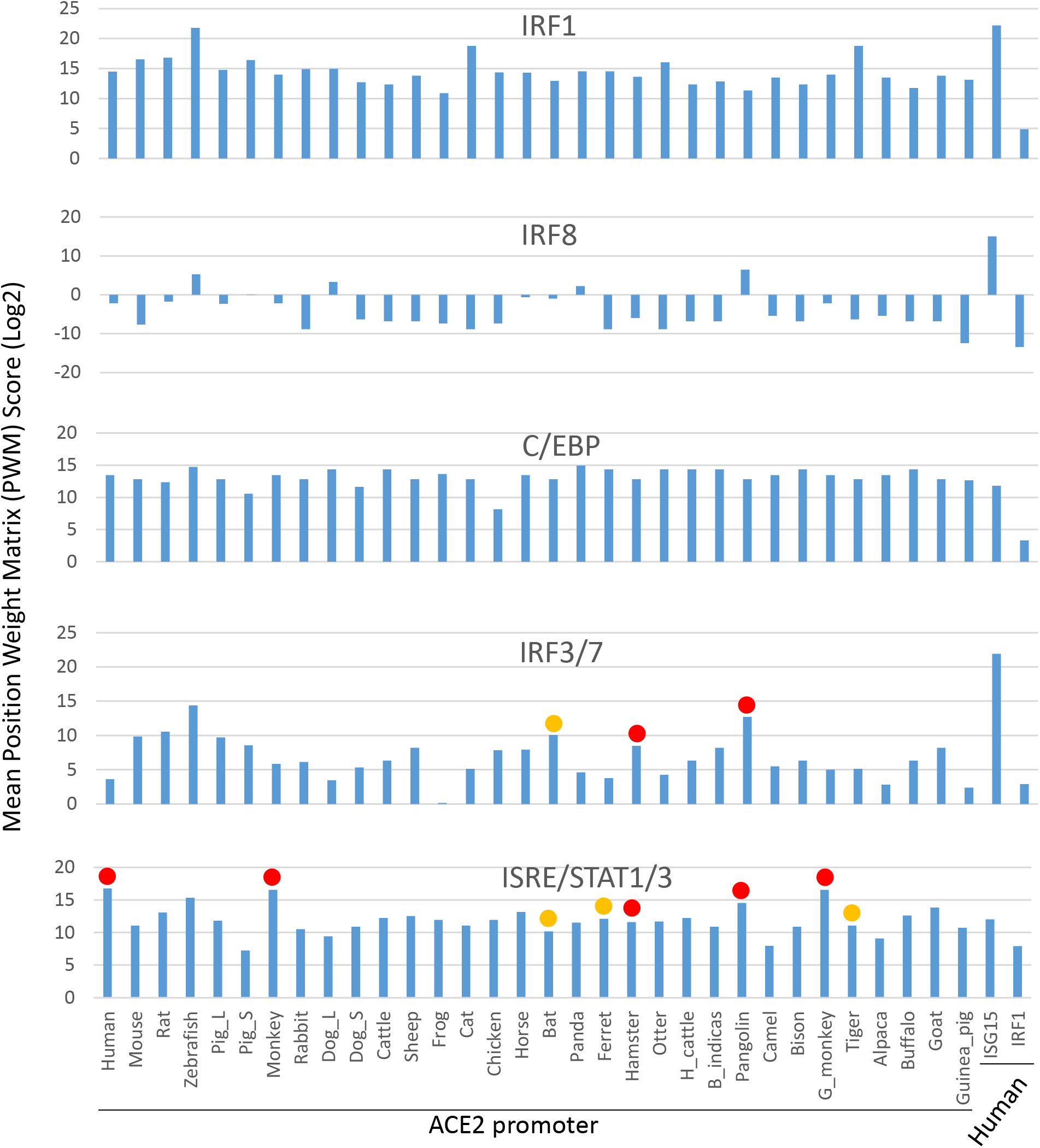
Scores of mean position weight matrix (PWM) of key *cis*-elements in the proximal promoters of *ACE2* genes that containing binding sites for canonical IFN-dependent transcription factors, which include ISRE/STAT1/3, IRF1, IRF3/7 and IRF8, as well as C/EBP as a core transcription factors for pro-inflammatory response. These IFN-dependent transcription factors, particularly IRF3/7 and ISRE/STAT critical for IFN stimulation, are differentially enriched in the promoter regions of *ACE2* genes in a species-dependent way. Especially, increased enrichment of ISRE/STAT1/3 and IRF3/7 binding sites are detected in the SARS-CoV2/COVID19 susceptible species (indicated with solid orange or red circles, respectively). In contrast, the PWM for IRF1 and C/EBP, which regulate inflammation, are less differentially enriched in ACE2 promoters from different animal species. The promoters of a typical human robust interferon-stimulated gene (ISG) 15 and IRF1 (a typical tunable ISG) are used as references. Higher enrichment of ISRE/STAT1/3 and IRF3/7 corresponds to SARS-CoV2 susceptibility in experimentally validated animal species and humans. Abbreviations: C/EBP, CCAAT/enhancer binding protein; IRF, interferon-regulatory factor; ISRE, Interferon-sensitive response element; STAT, Signal transducer and activator of transcription; PWM, position weight matrix. The PWM tools are used through https://ccg.epfl.ch/cgi-bin/pwmtools.

**Figure 7:**
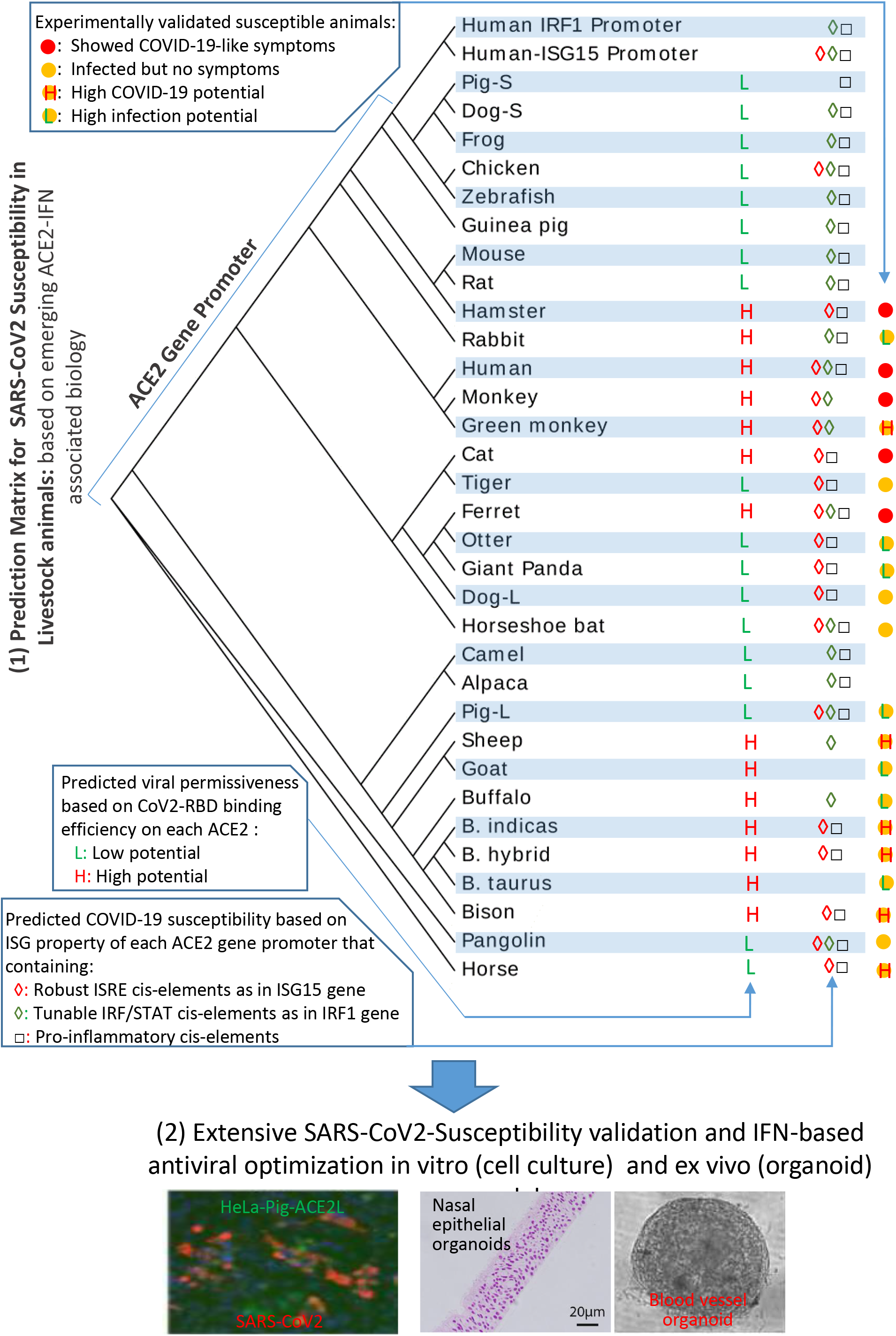
Prediction matrix and in vitro validation of SARS-CoV2 susceptibility in major vertebrate species based on (1) the binding capacity of the viral spike RBD on host ACE2 receptor; (2) the promotion of animal *ACE2* gene by the interferon response. Studies show that affinity adaption of the viral Spike receptor binding domain (RBD) and ACE2 receptor determines the viral susceptibility. SARS-CoV2 not only adapts a high binding affinity to human ACE2 for cell attachment, but also antagonizes the host antiviral interferon (IFN) response and utilizes the IFN-stimulated property of human *ACE2* gene to boost spreading.

### 3.6. Matrix scores of interferon-inductive elements in ACE2 *gene* promoters correspond to SARS-CoV2 susceptibility

The position weight matrix (PWM) stands as a position-specific scoring model for the binding specificity of a transcription factor (TF) on the DNA sequences [50]. Using PWM toolsets online (https://ccg.epfl.ch/cgi-bin/pwmtools), we evaluate mean PWM of key cis-elements in the proximal promoters of *ACE2* genes that containing binding sites for canonical IFN-dependent transcription factors, which include ISRE/STAT, IRF1. IRF3/7 and IRF8, as well as C/EBP representing a core transcription factor for pro-inflammation. These IFN-dependent transcription factors, particularly IRF3/7 and ISRE/STAT for IFN stimulation, are differentially enriched in the promoter regions of *ACE2* genes in a species-dependent way. Higher enrichment of ISRE/STAT1/3 and/or IRF3/7 binding sites are detected in most SARS-CoV2/COVID19 susceptible species (indicated with solid orange or red circles, respectively). In contrast, the PWM for IRF1 and C/EBP, which regulate inflammation, are less differentiated in ACE2 promoters from animal species, indicating that *ACE2* genes are more universally regulated by inflammation than that by the viral infection or IFN-induction in a species-dependent way (Fig. 6). As compared with the promoters of a typical human robust ISG15 and tunable IRF1 genes, this data indicate that *ACE2* genes (particularly the primate ones) are not typical robust or tunable ISGs as represented by ISG15 or IRF1, but respond differently to viral infection (through IRF3/7) or IFN auto-induction (via ISRE/STAT) in a species-dependent manner (Fig. 6) [48]. Higher enrichment of ISRE/STAT1/3 and/or IRF3/7 corresponds to SARS-CoV2 susceptibility in experimentally validated mammalian species especially primates, but not to the phylogenically distant species such as zebrafish, which has very low potential for SARS-CoV2 susceptibility due to the high disparity of ACE2 structures (Fig. 1 and Fig. S1). In addition, the proximal promoters of the pig and dog ACE2-S genes differ much in their IFN-responsive elements to most ACE2 promoters in mammalians (Fig. 5 and Fig. 6). However, they are phylogenically sister to the ACE2 promoters from the primitive vertebrates (frog, chicken and zebrafish) (Fig. 7, phylogenic tree). This indicates that the expression of these short ACE2 isoforms is more conservative than the long ACE2 isoforms, which represent a more recent evolution obtaining ACE2 epigenetic regulation by IFN-signaling (Fig. 7) [49].

### 3.7. Integrating ACE2 structural analysis, isoforms variants, and interferon-association for SARS-CoV2 susceptibility prediction and validation

Studies show that affinity adaption of the viral S-RBD and ACE2 receptor determines the cellular permissiveness to the virus [28,38,39]. SARS-CoV2 not only adapts a high binding affinity to human ACE2 for cell attachment, but also antagonizes host antiviral interferon (IFN) response and utilizes IFN-stimulated property of human *ACE2* gene to boost spreading [15,38,39,49]. In addition to structural analysis of simulated S-RBD-ACE2 interaction, we propose that several immunogenetic factors, including the evolution of S-binding-void ACE2 isoforms in some domestic animals, the species-specific IFN system, and epigenetic regulation of IFN-stimulated property of host *ACE2* genes, contribute to the viral susceptibility and the development of COVID-19-like symptoms in certain animal species [15,38,39,49]. A computational program in development that incorporates this multifactorial prediction matrix and in vitro validation of SARS-CoV2 susceptibility in major vertebrate species will be necessary to address public concerns relevant to SARS-CoV2 infections in animals (Fig. 7). It will also lead to the development of better animal models for anti-COVID19 investigations [21]. In addition, several IFN-based therapies for treatment of COVID19 have been proposed and are in the process of clinic trails [51–54]. Considering the viral stealth of IFN-stimulated property of human ACE2, a timely and subtype-optimized IFN treatment should be delivered rather than a general injection of typical human IFN-α/β subtypes [51–54]. In this line, domestic livestock like pigs and cattle have a most evolved IFN system containing numerous unconventional IFN subtypes. Some of these unconventional IFN subtypes, such as some porcine IFN-ω exert much higher antiviral activity than IFN-α even in human cells and most IFN-λ retain antiviral activity with less pro-inflammatory activity, could be utilized for developing effective antiviral therapies [55,56]. In summary, a predication matrix, which integrates the structural analysis of S-RBD-ACE2 interfacial interface and the species-specific immunogenetic diversity of *ACE2* genes, was used to predict the SARS-CoV2 susceptibility and fit current knowledge about the infectious potential already validated in different animal species (Fig. 7). More extensive validation experiments are needed to further improve this prediction matrix. Our current results demonstrate several previously unstudied immunogenetic properties of animal *ACE2* genes and imply some domestic animals, including dogs, pigs and cattle/goats, may obtain some immunogenetic diversity to confront SARS-CoV2 infection and face less COVID-19 risk than may have been previously thought. However, immediate biosecurity practices should be applied in animal management to reduce animal exposure to the virus and prevent potential species-specific adaptation (Fig. 7). For livestock breeding programs that targeting disease resistance to respiratory viruses, the genetic and epigenetic diversity of *ACE2* genes as well antiviral ISGs are highly recommended [48,49,55,56].

In conclusion, SARS-CoV2 evolves to fit well with human (and non-human primates) ACE2 receptor through the structural interfacial affinity, immunogenetic diversity and epigenetic expression regulation, which results in a highly infectious efficacy [1–3,15,28,38,39]. Most mammals, especially those that belong to glires, primates and carnivores, have a higher potential for SARS-CoV2 susceptibility but in a species-different manner based on the S-binding-void ACE2 isoforms and the difference of the IFN-inductive propensity of the major *ACE2* genes. Most ungulate animals appear have a low susceptibility potential with horses and sheep having a high potential (Fig. 7). Current development of IFN-based anti-COVID19 therapies should consider the ISG property of human *ACE2* gene to optimize for timely application using a highly-antiviral subtype that potentially have less anti-inflammatory activity [55–57]. The evolution of the IFN complex and functional diversity in domestic animals (such as pigs and cattle) provides a natural model for optimizing IFN antiviral regulation and therapy development [55,56].

## Supporting information

Supplemental Excel Sheet

## Author Contributions

E.R.S. and Y.T. contributed to idea conceptualization, data computation and draft preparation. Y.G. and L.C.M. help in conception and data discussion; L.C.M. helped for funding acquisition. Y.S. supervises overall conceptualization, data collection & process, computation, draft writing, and funding acquisition.

