## Supplemental Fig 1 for "Integrate Structural Analysis, Isoform Diversity, and Interferon-Inductive Propensity of ACE2 to Refine SARS-CoV2 Susceptibility Prediction in Vertebrates"

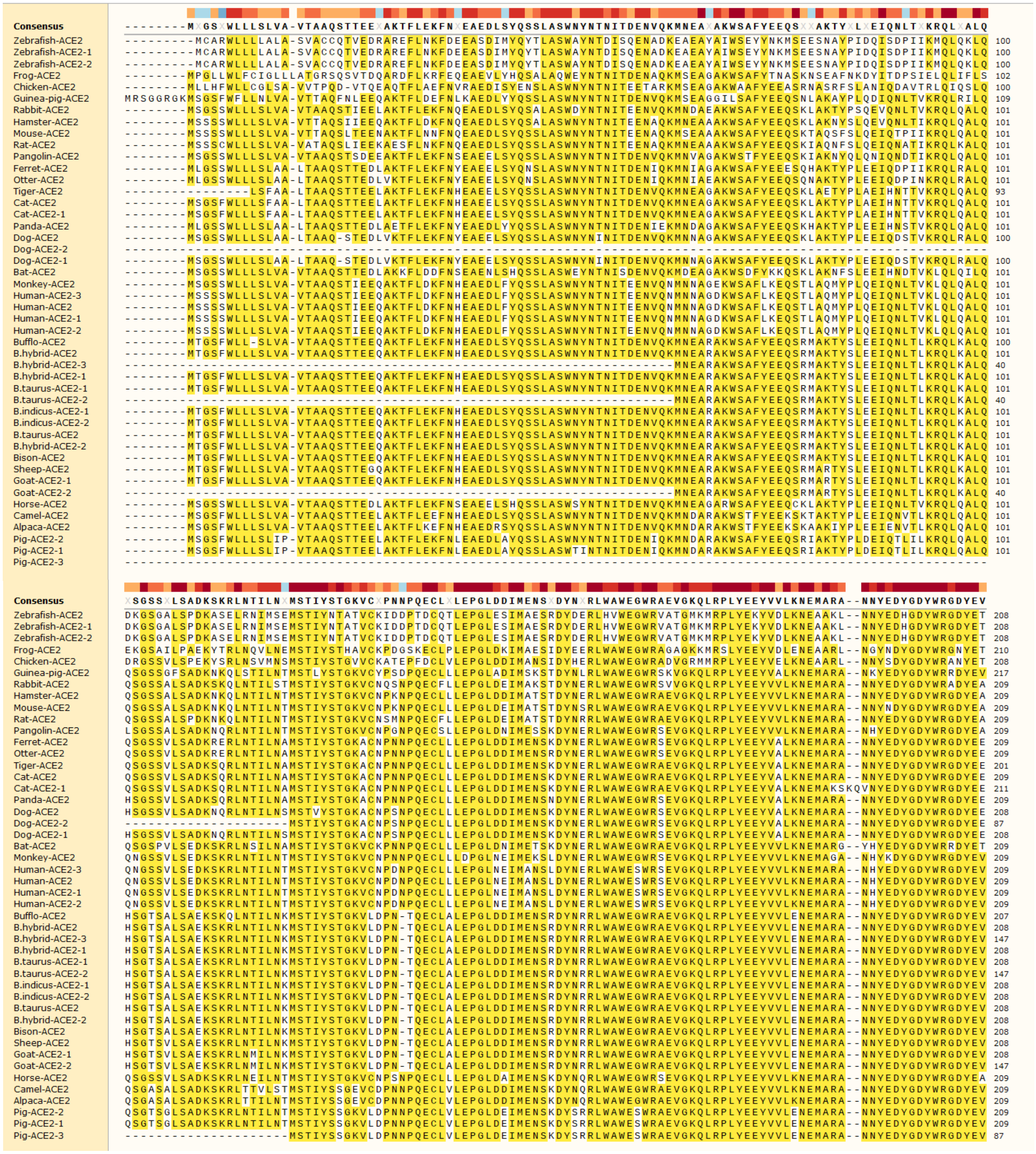
Consensus
Zebrafish-ACE2
Zebrafish-ACE2-1
Zebrafish-ACE2-2
Frog-ACE2
Chicken-ACE2
Guinea-pig-ACE2
Rabbit-ACE2
Hamster-ACE2
Mouse-ACE2
Rat-ACE2
Pangolin-ACE2
Ferret-ACE2
Otter-ACE2
Tiger-ACE2
Cat-ACE2
Cat-ACE2-1
Panda-ACE2
Dog-ACE2
Dog-ACE2-2
Dog-ACE2-1
Bat-ACE2
Monkey-ACE2
Human-ACE2-3
Human-ACE2
Human-ACE2-1
Human-ACE2-2
Bufflo-ACE2
B.hybrid-ACE2
B.hybrid-ACE2-3
B.hybrid-ACE2-1
B.taurus-ACE2-1
B.taurus-ACE2-2
B.indicus-ACE2-1
B.indicus-ACE2-2
B.taurus-ACE2
B.hybrid-ACE2-2
Bison-ACE2
Sheep-ACE2
Goat-ACE2-1
Goat-ACE2-2
Horse-ACE2
Camel-ACE2
Alpaca-ACE2
Pig-ACE2-2
Pig-ACE2-1
Pig-ACE2-3

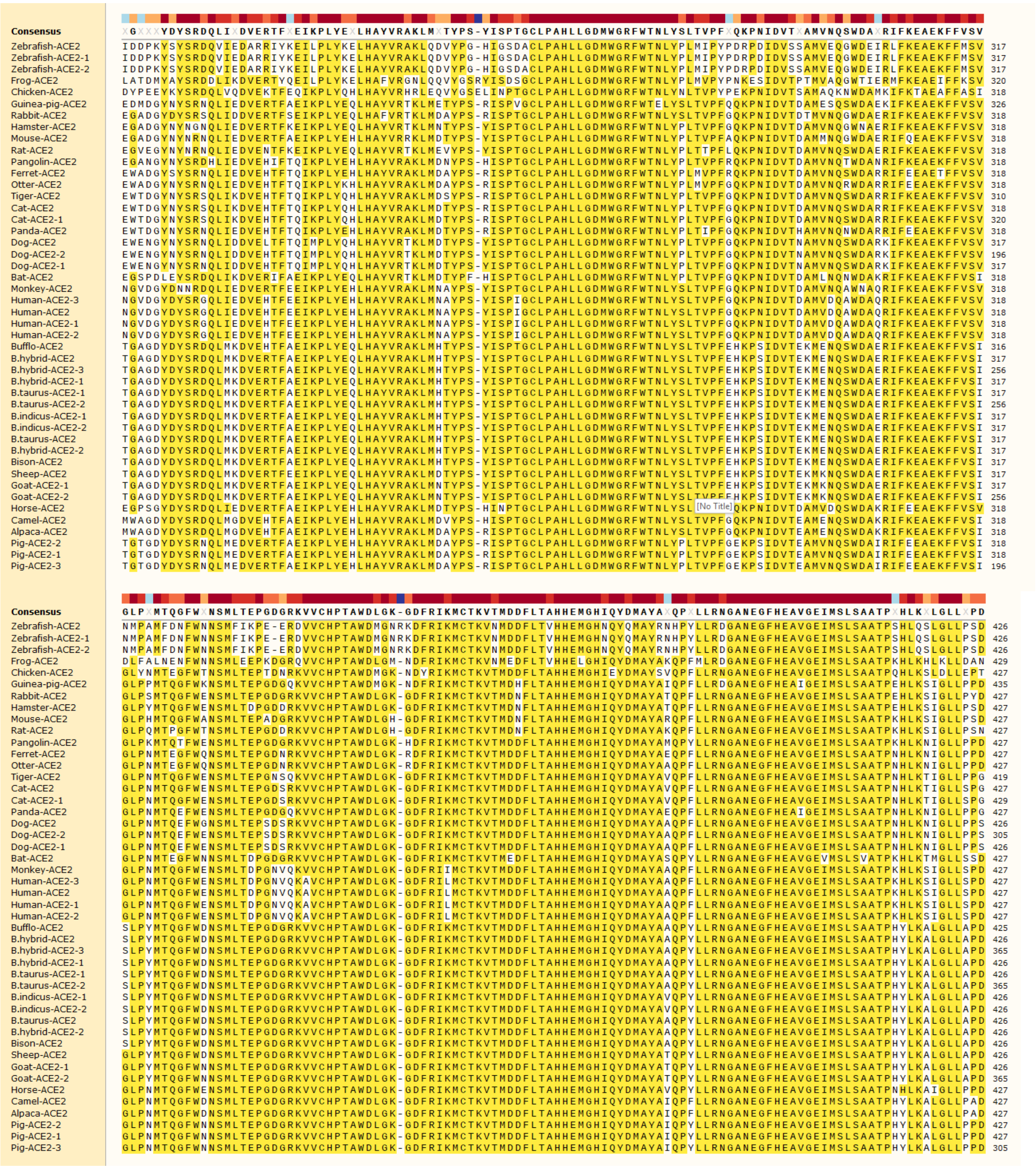
Consensus
XGXXXYDYSRDQLIXDVERTFXEIKPLYEXLHAYVRAKLMXTYPS-YISPTGCLPAHLLGDMWGRFWTNLYSLTVPFXQKPNIDVTXAMVNQSWDAXRIFKEAEKFFVSV
Zebrafish-ACE2
Zebrafish-ACE2-1
Zebrafish-ACE2-2
Frog-ACE2
Chicken-ACE2
Guinea-pig-ACE2
Rabbit-ACE2
Hamster-ACE2
Mouse-ACE2
Rat-ACE2
Pangolin-ACE2
Ferret-ACE2
Otter-ACE2
Tiger-ACE2
Cat-ACE2
Cat-ACE2-1
Panda-ACE2
Dog-ACE2
Dog-ACE2-2
Dog-ACE2-1
Bat-ACE2
Monkey-ACE2
Human-ACE2-3
Human-ACE2
Human-ACE2-1
Human-ACE2-2
Bufflo-ACE2
B.hybrid-ACE2
B.hybrid-ACE2-3
B.hybrid-ACE2-1
B.taurus-ACE2-1
B.taurus-ACE2-2
B.indicus-ACE2-1
B.indicus-ACE2-2
B.taurus-ACE2
B.hybrid-ACE2-2
Bison-ACE2
Sheep-ACE2
Goat-ACE2-1
Goat-ACE2-2
Horse-ACE2
Camel-ACE2
Alpaca-ACE2
Pig-ACE2-2
Pig-ACE2-1
Pig-ACE2-3
[No Title]
Consensus
GLPXMTQGFWXNSMLTEPGDGRKVVCHPTAWDLGK-GDFRIKMCTKVTMDDFLTAHHEMGHIQYDMAYAXQPXLLRNGANEGFHEAVGEIMSLSAATPXHLKXLGLLXPD

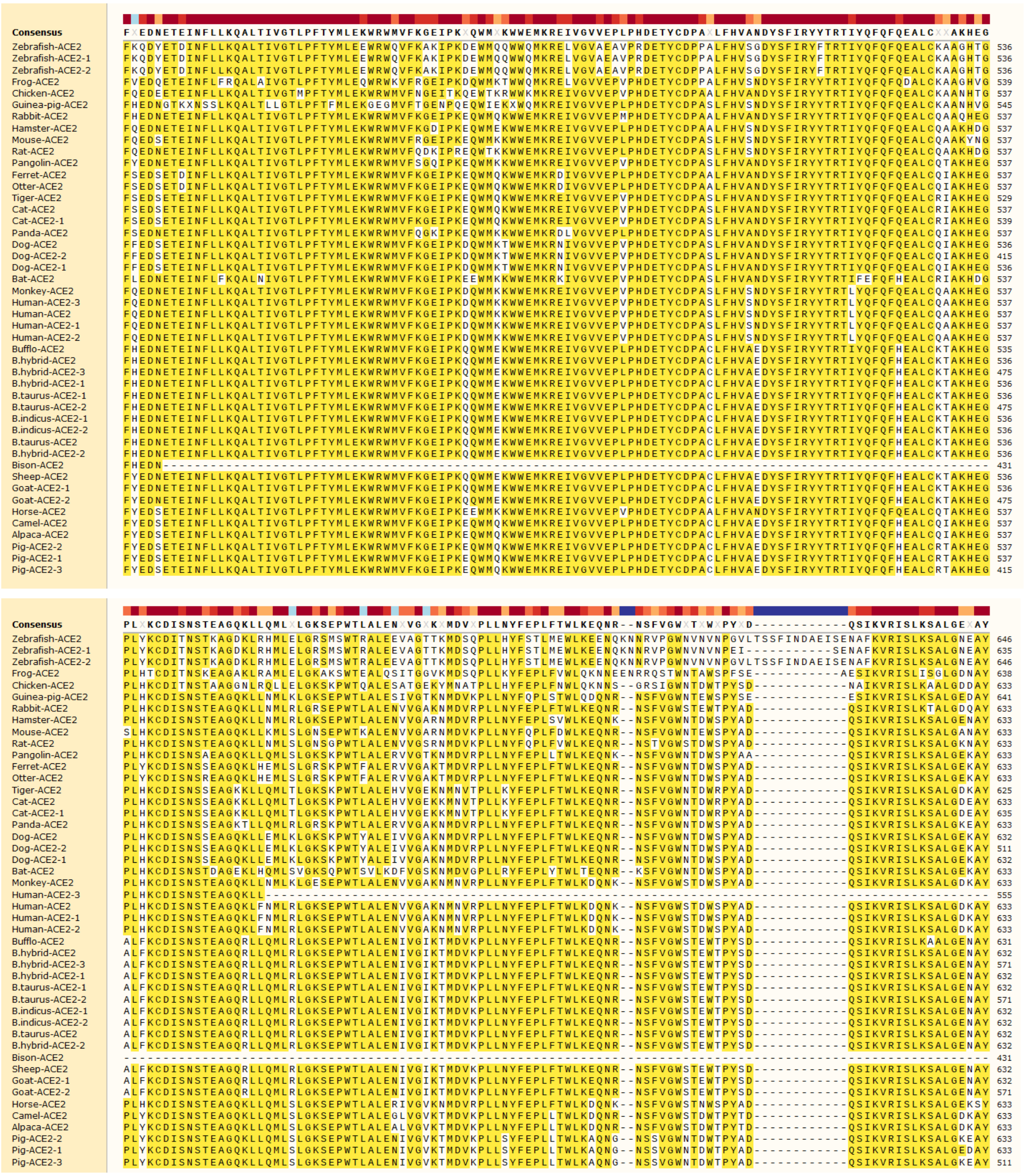
Consensus
Zebrafish-ACE2
Zebrafish-ACE2-1
Zebrafish-ACE2-2
Frog-ACE2
Chicken-ACE2
Guinea-pig-ACE2
Rabbit-ACE2
Hamster-ACE2
Mouse-ACE2
Rat-ACE2
Pangolin-ACE2
Ferret-ACE2
Otter-ACE2
Tiger-ACE2
Cat-ACE2
Cat-ACE2-1
Panda-ACE2
Dog-ACE2
Dog-ACE2-2
Dog-ACE2-1
Bat-ACE2
Monkey-ACE2
Human-ACE2-3
Human-ACE2
Human-ACE2-1
Human-ACE2-2
Bufflo-ACE2
B.hybrid-ACE2
B.hybrid-ACE2-3
B.hybrid-ACE2-1
B.taurus-ACE2-1
B.taurus-ACE2-2
B.indicus-ACE2-1
B.indicus-ACE2-2
B.taurus-ACE2
B.hybrid-ACE2-2
Bison-ACE2
Sheep-ACE2
Goat-ACE2-1
Goat-ACE2-2
Horse-ACE2
Camel-ACE2
Alpaca-ACE2
Pig-ACE2-2
Pig-ACE2-1
Pig-ACE2-3

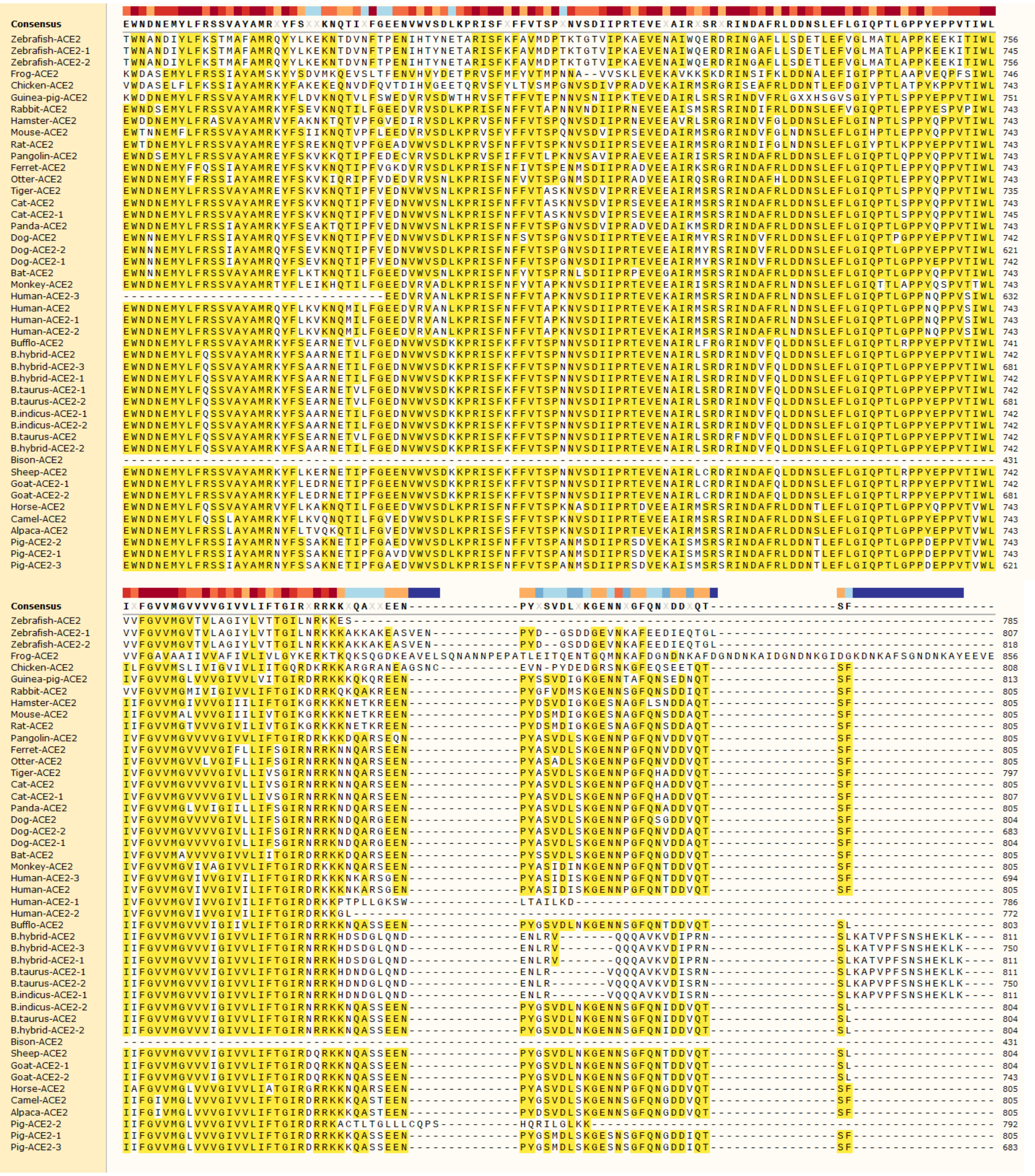
Consensus
EWNDNEMYLFRSSVAYAMR x YFS xx KNQTI x FGEENVWVSDLKPRISF x FFVTSP x NVSDIIPRTEVE x AIR x SR x RINDAFRLDDNSLEFLGIQPTLGPPYEPPVTIWL
Zebrafish-ACE2
Zebrafish-ACE2-1
Zebrafish-ACE2-2
Frog-ACE2
Chicken-ACE2
Guinea-pig-ACE2
Rabbit-ACE2
Hamster-ACE2
Mouse-ACE2
Rat-ACE2
Pangolin-ACE2
Ferret-ACE2
Otter-ACE2
Tiger-ACE2
Cat-ACE2
Cat-ACE2-1
Panda-ACE2
Dog-ACE2
Dog-ACE2-2
Dog-ACE2-1
Bat-ACE2
Monkey-ACE2
Human-ACE2-3
Human-ACE2
Human-ACE2-1
Human-ACE2-2
Bufflo-ACE2
B.hybrid-ACE2
B.hybrid-ACE2-3
B.hybrid-ACE2-1
B.taurus-ACE2-1
B.taurus-ACE2-2
B.indicus-ACE2-1
B.indicus-ACE2-2
B.taurus-ACE2
B.hybrid-ACE2-2
Bison-ACE2
Sheep-ACE2
Goat-ACE2-1
Goat-ACE2-2
Horse-ACE2
Camel-ACE2
Alpaca-ACE2
Pig-ACE2-2
Pig-ACE2-1
Pig-ACE2-3
Consensus
I x FGVVMGVVVVGIVVLIFTGIR x RRKK x QA xx EEN-------------PY x SVDL x KGENN x GFQN x DD x QT---------------SF-----------------

| | | |
|---|---|---|
| **Consensus** | ------ |  |
| Zebrafish-ACE2 | ------ | 785 |
| Zebrafish-ACE2-1 | ------ | 807 |
| Zebrafish-ACE2-2 | ------ | 818 |
| Frog-ACE2 | TMNTTF | 862 |
| Chicken-ACE2 | ------ | 808 |
| Guinea-pig-ACE2 | ------ | 813 |
| Rabbit-ACE2 | ------ | 805 |
| Hamster-ACE2 | ------ | 805 |
| Mouse-ACE2 | ------ | 805 |
| Rat-ACE2 | ------ | 805 |
| Pangolin-ACE2 | ------ | 805 |
| Ferret-ACE2 | ------ | 805 |
| Otter-ACE2 | ------ | 805 |
| Tiger-ACE2 | ------ | 797 |
| Cat-ACE2 | ------ | 805 |
| Cat-ACE2-1 | ------ | 807 |
| Panda-ACE2 | ------ | 805 |
| Dog-ACE2 | ------ | 804 |
| Dog-ACE2-2 | ------ | 683 |
| Dog-ACE2-1 | ------ | 804 |
| Bat-ACE2 | ------ | 805 |
| Monkey-ACE2 | ------ | 805 |
| Human-ACE2-3 | ------ | 694 |
| Human-ACE2 | ------ | 805 |
| Human-ACE2-1 | ------ | 786 |
| Human-ACE2-2 | ------ | 772 |
| Bufflo-ACE2 | ------ | 803 |
| B.hybrid-ACE2 | ------ | 811 |
| B.hybrid-ACE2-3 | ------ | 750 |
| B.hybrid-ACE2-1 | ------ | 811 |
| B.taurus-ACE2-1 | ------ | 811 |
| B.taurus-ACE2-2 | ------ | 750 |
| B.indicus-ACE2-1 | ------ | 811 |
| B.indicus-ACE2-2 | ------ | 804 |
| B.taurus-ACE2 | ------ | 804 |
| B.hybrid-ACE2-2 | ------ | 804 |
| Bison-ACE2 | ------ | 431 |
| Sheep-ACE2 | ------ | 804 |
| Goat-ACE2-1 | ------ | 804 |
| Goat-ACE2-2 | ------ | 743 |
| Horse-ACE2 | ------ | 805 |
| Camel-ACE2 | ------ | 805 |
| Alpaca-ACE2 | ------ | 805 |
| Pig-ACE2-2 | ------ | 792 |
| Pig-ACE2-1 | ------ | 805 |
| Pig-ACE2-3 | ------ | 683 |

Supplemental Fig. S1: Multiple sequence alignment of vertebrate ACE2 orthologs/paralogs. Sequence of ACE2 protein precursors were extracted from NCBI Gene and relevant databases (https://www.ncbi.nlm.nih.gov/gene). The protein sequences were aligned using the multiple sequence alignment tools of ClustalW or Muscle through an EMBL-EBI port (https://www.ebi.ac.uk/) under the default parameters.
